# Teaching free energy calculations to learn from experimental data

**DOI:** 10.1101/2021.08.24.457513

**Authors:** Marcus Wieder, Josh Fass, John D. Chodera

## Abstract

Alchemical free energy calculations are an important tool in the computational chemistry toolbox, enabling the efficient calculation of quantities critical for drug discovery such as ligand binding affinities, selectivities, and partition coefficients. However, modern alchemical free energy calculations suffer from three significant limitations: (1) modern molecular mechanics force fields are limited in their ability to model complex molecular interactions, (2) classical force fields are unable to treat phenomena that involve rearrangements of chemical bonds, and (3) these calculations are unable to easily learn to improve their performance if readily-available experimental data is available. Here, we show how all three limitations can be overcome through the use of quantum machine learning (QML) potentials capable of accurately modeling quantum chemical energetics even when chemical bonds are made and broken. Because these potentials are based on mathematically convenient deep learning architectures instead of traditional quantum chemical formulations, QML simulations can be run at a fraction of the cost of quantum chemical simulations using modern graphics processing units (GPUs) and machine learning frameworks. We demonstrate that alchemical free energy calculations in explicit solvent are especially simple to implement using QML potentials because these potentials lack singularities and other pathologies typical of molecular mechanics potentials, and that alchemical free energy calculations are highly effective even when bonds are broken or made. Finally, we show how a limited number of experimental free energy measurements can be used to significantly improve the accuracy of computed free energies for unrelated compounds with no significant generalization gap. We illustrate these concepts on the prediction of aqueous tautomer free energies (related to tautomer ratios), which are highly relevant to drug discovery in that more than a quarter of all approved drugs exist as a mixture of tautomers.

## Introduction

Alchemical free energy calculations are a versatile tool for estimating free energy differences between physical systems. Relative free energy calculations, in particular, enable the efficient computation of free energies of transfer—such as ligand binding free energies or octanol-water partition coefficients—between related molecules, making them one of the most useful tools in the computational chemistry toolbox [1].

While alchemical free energy calculations have been used extensively in structure-enabled drug discovery [2–4], current generation molecular mechanics alchemical free energy calculations have several limitations that severely constrain their domain of applicability [5]: First, they rely on molecular mechanics force fields which are generally fit to unrelated quantum chemical and experimental data, making compromises that may limit accuracy due to the limited flexibility of the potential functional forms [6, 7]. While bespoke fitting tools that derive some or all parameters from quantum chemistry [8–12] or fast quantum machine learning surrogates [13] have recently become popular, these approaches are incapable of learning from experimental data collected on related molecules. Second, the functional forms for popular class I molecular mechanics force fields (such as GAFF [14, 15], CGenFF [16], or OPLS [11, 17]) and even class II force fields [6,7] are highly limited, omitting high-order terms and important couplings—such as torsion-torsion couplings now deemed essential for accurate peptide backbone energetics [18–20] but omitted from small molecule force fields—leading to significant deviations from quantum chemical energetics [21]. Third, molecular mechanics force fields are incapable of modeling phenomena that involve the rearrangements of chemical bonds, since their functional forms do not permit the formation or breaking of chemical bonds [6, 7].

Here, we demonstrate that alchemical free energy calculations reformulated for a new generation of quantum machine learning (QML) potentials [22] are capable of addressing all of these issues. We consider a challenging system that typifies all three of these challenges: The calculation of tautomer ratios for drug-like organic molecules. While over a quarter of approved drugs significantly populate multiple tautomers in solution [23], these quantities cannot be computed using classical force fields because they involve re-arrangements of chemical bonds, and the accuracy of relative free energies of hydration computed with traditional molecular mechanics force fields cannot systematically be improved. As we shall demonstrate, not only are alchemical free energy calculations *easier* to perform with QML potentials, but they can also give accuracy close to quantum chemical energetics that can systematically be improved using a limited quantity of experimental data.

### Quantum machine learning potentials present new opportunities for alchemical free energy calculations

Recently, accurate and transferable quantum machine learning(QML) potentials (e.g. ANI [24], PhysNet [25], and SchNet [26]) aiming to reproduce the energetics of high-level quantum nuclear potentials (within the Born-Oppenheimer approximation) by leveraging modern machine learning architectures have been successfully applied to model organic small molecules. QML potentials can be used as a substitute for quantum mechanical treatment of organic molecular systems and—given sufficient and appropriate training data—are able to reproduce quantum chemical energies and forces without significant loss of accuracy but with orders of magnitude less computational cost than the quantum chemical methods they aim to reproduce. For example, the ANI-1x neural network potential is able to reproduce DFT-level energies (calculated with the *ω*B97X functional and the 6-31G* basis set) six orders of magnitude faster with an inaccuracy of only 1 kcal/mol [24]; second-generation QML potentials, such as ANI-1ccx, leverage transfer learning to approach extreme CCSD(T)/CBS accuracy with essentially no additional computational cost [27].

In this work, we show how QML potentials (here, ANI-2x [28]) can be modified to perform alchemical relative free energy calculations in explicitly solvated systems where the entire system (here, a droplet) is treated with the quantum machine learning potential. We demonstrate how this approach can be used to compute aqueous tautomer ratios—one of the most challenging problems for free energy calculations, since its success depends on both a highly accurate estimate of both the quantum chemical energy for breaking and making covalent bonds as well as the relative hydration or solvation free energy of two distinct chemical species [29]. Systematic investigations in tautomer ratios in the literature are scarce, and their findings are often inconclusive or difficult to generalize [23, 30–32]. We hope to add significantly to the canon of available studies and data about tautomer ratios with a systematic investigation of 354 tautomer pairs, as well as demonstrate how alchemical free energies with QML potentials are poised to significantly extend the domain of applicability, accuracy, and systematic improvability of alchemical free energy calculations.

### QML potentials can be trained using experimental thermodynamic data to systematically improve accuracy

QML potentials—such as those from the ANI family of potentials [24, 28, 33]—offer significant advantages over both molecular mechanics potentials (due to their accuracy) and quantum chemical calculations (due to their speed), but there is third significant advantage that has not yet been exploited: Because QML models are generally implemented within auto-differentiation capable frameworks (such as PyTorch [34], TensorFlow [35], or JAX [36]), the entire calculation becomes differentiable, enabling systematic improvement of the accuracy of these potentials by retraining to experimental free energies or ensemble averages.

To *efficiently* optimize an ensemble property (such as an expectation or free energy difference) to fit experimental observables, it is necessary to formulate a framework in which these properties and their gradients with respect to parameters can be extrapolated to new parameter values without requiring the entire simulation be performed each time the parameter is perturbed. Traditional approaches to computing these parameter gradients have relied on *sensitivity analysis* (evaluating the partial derivatives of a simulation average with respect to simulation parameters during a simulation), using analytical expressions for the derivatives [37, 38]. To test the effect of the new parameter values on an ensemble property, simulations would then have to be performed with the new parameter set, making the optimization cycle (*training epoch*, in machine learning terms) both slow and computationally expensive [37, 38].

Here, we present a different approach based on simple gradients of the free energy and differentiable importance sampling. As starting point, we use the ANI QML potential with the published parameter set [28], which was derived by fitting gas phase quantum chemical energies and forces, and focus on systematically improving the parameters to predict accurate free energy differences in the condensed phase. We show how a modest-sized set (212 tautomer pairs) of experimentally determined tautomer ratios can be used as training data to refit the QML model parameters while adding a regularization term to minimize the required perturbations to quantum chemical energetics. To do this efficiently, we perform alchemical sampling and free energy calculations with the original parameter set (shown as **Alchemical sampling** and **Free energy estimate** phases in Figure 3 **A** and **B**). The loss function is then defined as the squared error between calculated and experimental tautomeric free energies, which is fully differentiable with respect to the QML potential neural net parameters. By calculating the parameter gradient of the loss and updating the parameters accordingly, a perturbed tautomeric free energy can be calculated by importance sampling. Practically, this is done via a reweighting scheme that estimates the free energy difference and its gradient with respect to the QML potential parameters from already-sampled configurations [39, 40]. In particular, we make use of the following relationship for any observable 〈*O*〉 for a target distribution *p* where 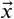 are samples from a different distribution *q*

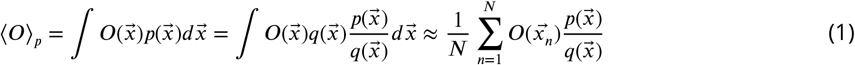

Only the numerator of the sampling weights 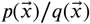 is modified during the **Parameter optimization via gradient descent** phases in Figure 3 since 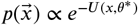 for the new parameter set *θ*\*, whereas the previous unnormalized weights 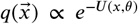 and sampled configurations 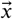 remain fixed during the optimization process. Provided changes to the original parameter set are small, importance sampling will have tolerable variance for the perturbed free energies (shown in Figure S.I.5). The gradient calculation, parameter update, and calculation of a new perturbed, differentiable free energy estimate is performed repeatedly until a stop condition is met.

Since alchemical sampling (the time consuming step) is performed once and conformations and energies are stored, this approach is highly efficient. Generating sufficient samples to estimate the tautomeric free energy difference for a single tautomer pair takes roughly 500 CPU hours using current software frameworks (11 sampled states × 48 CPU hours per state), though continual optimizations in both software and hardware are rapidly reducing the cost of QML calculations. The sampling step can be run in parallel on 11 CPUs for a single tautomer pair, each for 48h. Calculating the perturbed free energy, calculating the gradient, and performing an optimization step for a single tautomer pair takes just minutes, making the computation of gradients and parameter updates for a batch of the training set possible in 2-3 CPU hours, depending on the number of snapshots per intermediate state used for the free energy estimate. The full workflow is illustrated in Figure 3 and described in detail in the Methods section.

This easily-implemented approach offers a new paradigm for one of the more challenging tasks in biophysics: the development of small molecule potentials that can reproduce thermodynamic observables from ensemble data (e.g.[7, 41]). While this approach is not new, recent advances and increase interest in the field of QML potentials have motivated new developments in this field (e.g., [42–44]).

## QML potentials enable the efficient and accurate calculation of free energies by using QM based Hamiltonian

We investigated a subset of 354 experimentally well-determined tautomer ratios (selected via criteria described in **Methods**) obtained from the publicly available *Tautobase* [45]. We performed relative alchemical free energy calculations of tautomer pairs in solution using both ANI-1ccx and ANI-2x. The system was modeled using a water droplet surrounding the small molecule to mimic first-shell solvent effects without incurring the currently prohibitive expense of modeling fully QML solvent boxes. For a typical tautomer simulation, this resulted in a molecular system with approximately 220 to 260 atoms—one such system is shown in Figure 1 **A**.

**Figure 1.**
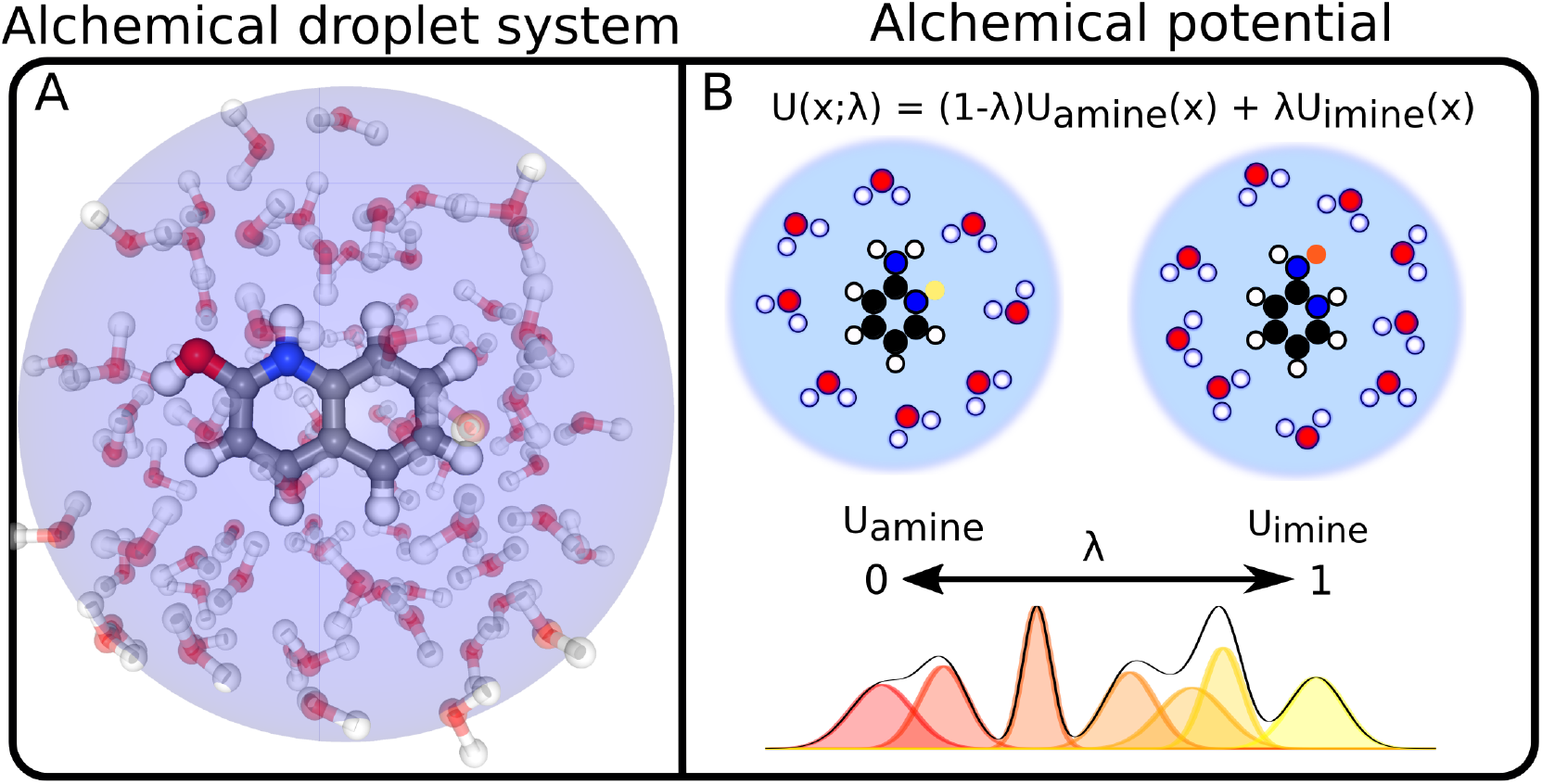
Quantum machine learning potentials can be used to rigorously calculate tautomeric free energy differences in solution. A single topology approach was used in this work, where real atoms are transmuted into no-interacting dummy atoms (restrained to remain in their chemically relevant position), and vice-versa. Panel **A** shows a snapshot of the topology of a tautomer system at *λ* = 0.5. The water droplet (16 Angstrom (Å) diameter) is indicated by the blue sphere. Panel **B** shows how linearly interpolating between the potential energy function at the two physical endstates (i.e. *U*_amine_ and *U*_imine_) generates the potential energy function at intermediate states (*λ* is used as control parameter). Dummy atoms are indicated by the yellow highlighted hydrogen at *U*_amine_ and the orange highlighted atom at *U*_imine_. The endstate and bridging distributions are used to estimates the tautomeric free energy difference in solution.

Alchemical free energy calculations were performed using a single-topology approach with 11 alchemical states by linearly interpolating between the endpoint potentials with a parameter *λ* ∈ [0,1] (Figure 1 **B**). Here, the QML potential includes the full system of small molecule and water molecules, with a harmonic potential applied to the water molecules to ensure they remain within the droplet. To avoid bond breaking or formation in each tautomeric endstate, chemical identity was enforced with flat bottom distance restraints applied to all bonds, with the minimum distance at which the restraint becomes nonzero selected so as not to interfere with normal bond stretch fluctuations. Details of the approach are given in **Methods**.

Given this experimental setting, we can ask two important questions: using the same tautomer dataset and simulation protocol, (1) is the predictive accuracy sensitive to the presence of explicit solvent, and (2) does using a QML model fit to a higher level of QML theory systematically improve tautomer ratio prediction accuracy? To obtain accurate predictions of tautomeric free energy differences, a highly accurate estimate of both the intrinsic free energy difference between tautomer pairs as well as solvation effects is needed [46]. It seems intuitive that the answer to both questions posed should be a resounding “yes”—yet, finding these trends in computational chemistry is often elusive and difficult (see, for example, [47**?**]).

### Does adding explicit solvent improve the accuracy of tautomer ratio predictions?

To assess whether the added complexity and computational cost of carrying out an alchemical free energy calculation to predict tautomer ratios in a water droplet led to an improvement in the predicted tautomer ratios, we compared our computed free energies using ANI-1ccx to our previous work using the same potential and method in vacuum [48]. These results can be directly compared since the same set of molecules was used to calculate the tautomeric free energies. The inclusion of a solvent droplet reduces the predicted tautomer free energy RMSE from 6.7 [5.8,7.7] kcal/mol to 4.8 [4.4,5.3] kcal/mol—an improvement of 2 kcal/mol—indicating that the addition of first-shell water interactions significantly improves the accuracy of tautomeric free energy estimates.

### Does improving quantum chemical level of theory improve tautomer ratio prediction accuracy?

Previous studies proposed the minimum level of theory for the gas-phase electronic energy to obtain tautomer ratios with an error of less than 1.5 kcal/mol was suggested to be MP2/pVDZ [29]. This suggestion was a result of the SAMPL2 blind predictive tautomer prediction challenge and—given the small set of tautomer pairs used in the challenge—should be considered with caution, as it also ignores the approximations introduced by the calculation of the standard state free energy at finite temperature. Intuitively, it would seem sensible that a higher level of QM theory should lead to less deviation from the true, experimental values. While ANI-2x was trained on energies/forces calculated with *ω*B97x/6-31G*(d) (below the level of theory suggested to be necessary for accurate tautomer ratio predictions), ANI-1ccx was retrained via transfer learning to energies recalculated with coupled cluster QM calculations considering single, double, and perturbative triple excitations (CCSD(T)) with an extrapolation to the complete basis set limit (CBS) [33]. CCSD(T)/CBS is the gold standard for electronic energy prediction and is able to predict thermochemical properties to within the limit of chemical accuracy (e.g., [49]).

We find that tautomeric free energy estimates obtained with ANI-2x (trained on the lower level of theory, *ω*B97x/6-31G*(d)) had an RMSE to experiment of 7.4 [95% CI: 6.8,7.8] kcal/mol. For ANI-1ccx, retrained to the gold standard CCSD(T)/CBS level of theory, the RMSE was indeed considerably lower, 4.8 [95%CI:4.4,5.3] kcal/mol (both alchemical free energy results are shown in Figure 2). These results indicate that the level of theory used to train ANI-2x is insufficient to accurately calculate tautomeric free energies, even if rigorous free energy calculations in explicit solvent are performed (and conformational degrees of freedom are sampled explicitly, in contrast to *e.g*. the rigid-rotor harmonic oscillator approximation).

**Figure 2.**
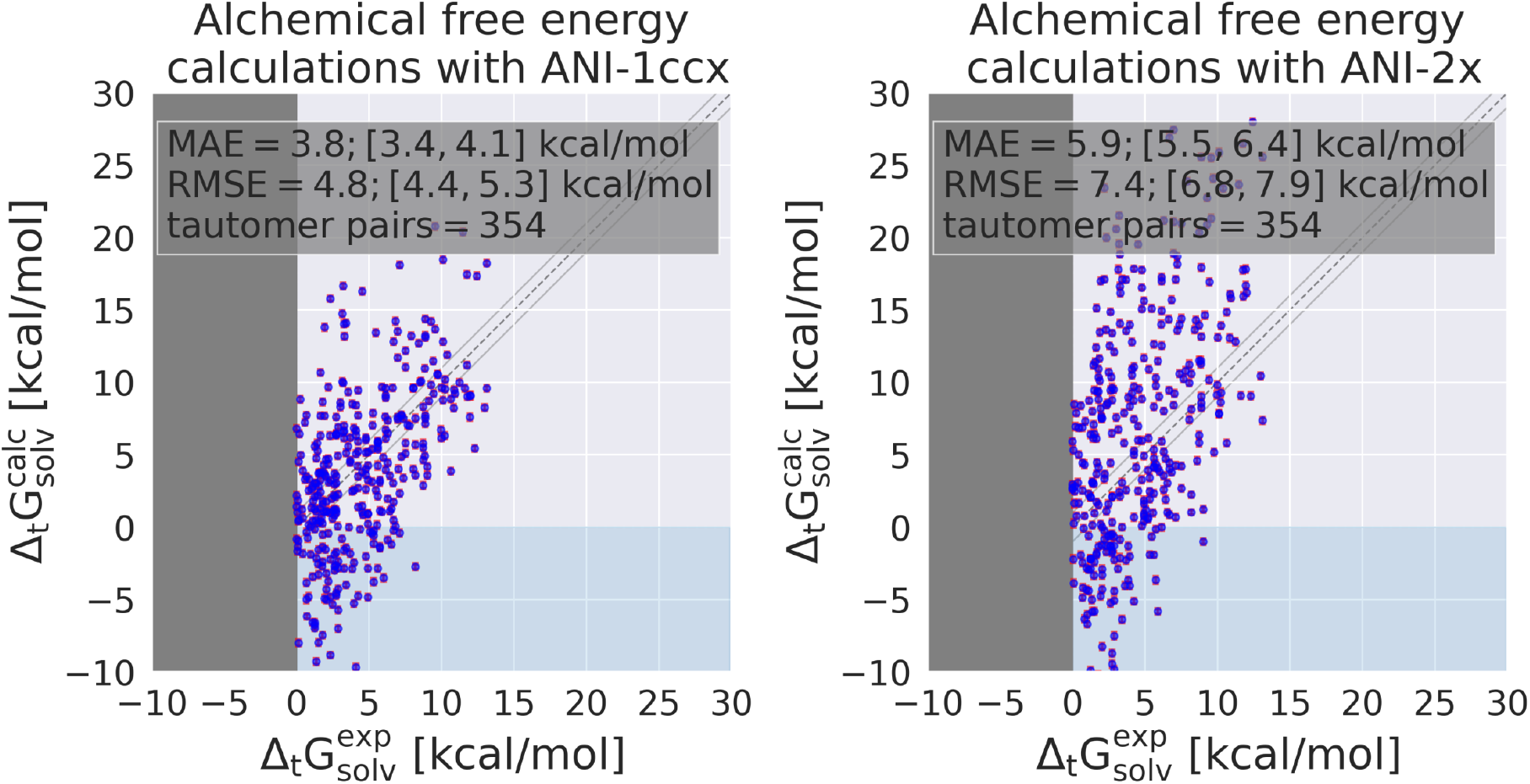
Alchemical relative free energy calculations using default ANI-2x parameters [33] have a large RMSE of 7.4 kcal/mol on the used subset of the Tautobase [45]. Error bars representing the standard error are shown in red for each of the 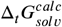 estimates.

Surprisingly, however, despite the significantly improved accuracy of ANI-1ccx compared to ANI-2x, ANI-1ccx proved to have a significant and critical defect that rendered it generally unsuitable for simulations of solvated systems. In particular we found that water molecules adopted highly nonphysical minimum geometries during the simulations that contributed very favorable to the total energy of the system and unfavorable to the ensemble standard deviation of the energy calculation but found ourselves unable to control this issue by penalizing increased ensemble standard deviation or/and deviating angles. We discuss these issues in detail in the **Analysis of ANI-1ccx droplet simulation** section in the Supplementary Information. We therefore focused exclusively on ANI-2x for the remainder of this study.

## Retraining a QML potential on a small set of experimental free energies systematically improves the accuracy of alchemical free energy calculations

Expressing free energy estimates as differentiable functions of force field parameters (or, in the case of QML, neural net parameters) allows the use of gradient-based optimization techniques to improve the accuracy of computed free energies in reproducing experimental data. While this would normally require expensive new alchemical simulations after every parameter update, we have developed a workflow that uses information-efficient reweighting estimators [40] to iteratively optimize the ANI-2x model to obtain improved free energy estimates without the need to re-run simulations, as illustrated in Figure 3.

**Figure 3.**
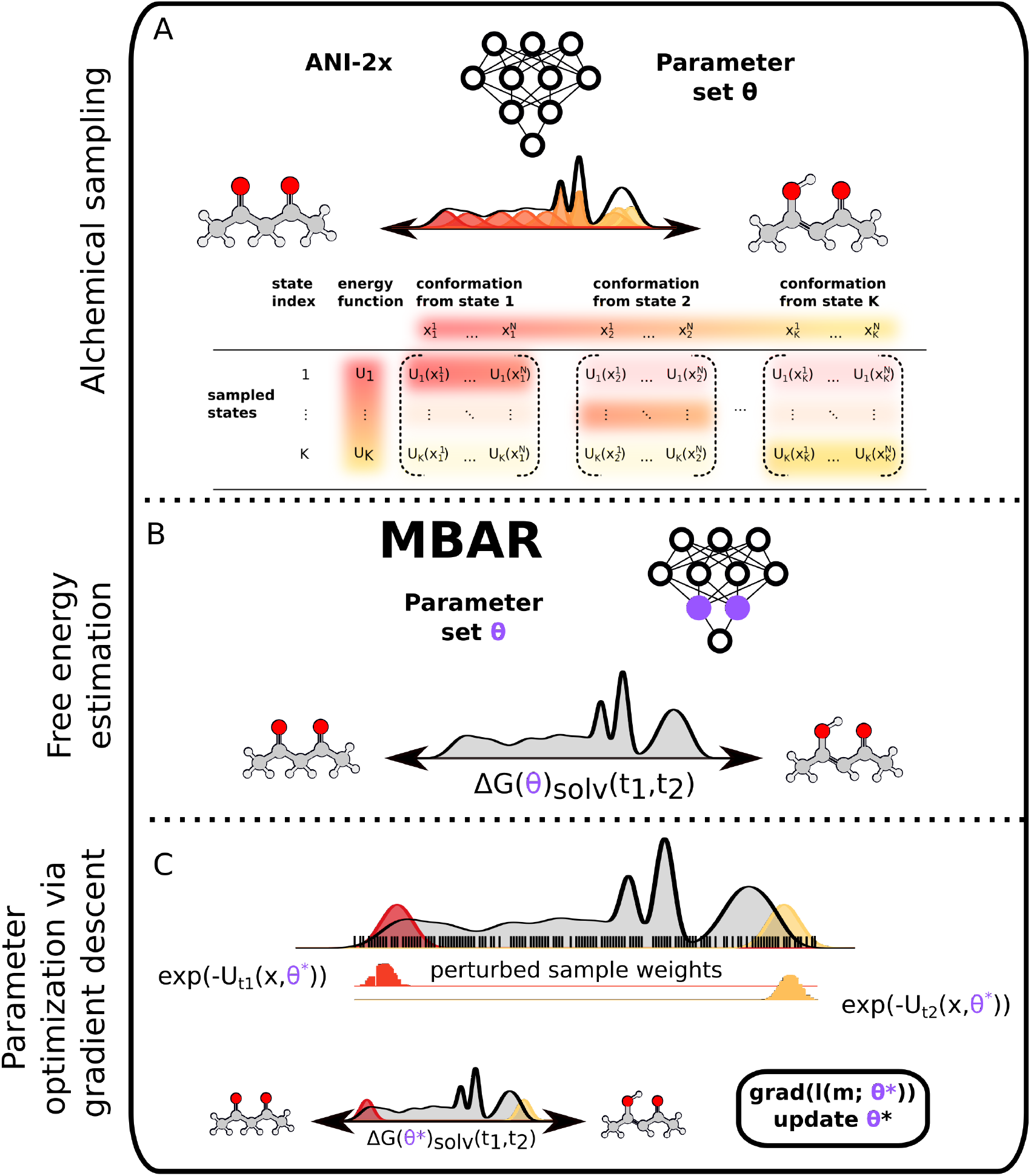
Reweighting techniques enable QML potentials to be efficiently optimized without new simulations. The optimization workflow can be split in three phases: **A** shows how conformations 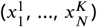 are sampled from the physical endstates and the alchemical bridging distributions using the original ANI-2x parameter set (*θ*) for the *K* sampled states and the *N* number of snapshots per state. Here, 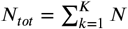, with *N* = 200 snapshots per alchemical state (*λ_k_*). These samples are then reevaluated using the energy functions (*U*_1_,…, *U_K_*) for all *K* alchemical states to construct an MBAR estimator for free energy differences between the physical tautomeric states [40]. **B** shows how the free energy estimate can be obtained for the native ANI-2x parameter set *θ*. **A** and **B** are the time consuming steps in this pipeline, consuming around 500 CPU hours (*K* = 11 alchemical *λ* states × 48 CPU hours per alchemical state) to sample the *K* alchemical states and estimate the tautomeric free energy difference. Steps **A** and **B** need be carried out once, and the snapshots obtained reused to estimate the tautomeric free energy difference and its parameter gradient for new QML potential neural net parameters *θ*\* near *θ*. **C** describes the parameter optimization workflow, where we optimize the QML potential parameters *θ* for only the last layer of the ensemble of ANI-2x atomic neural net parameters, retaining the earlier layers that perceive quantum chemical features. To obtain a perturbed free energy estimate, the potential energy and parameter gradient for each snapshot is recomputed using the perturbed parameter set (*θ*\*) and the new tautomer free energy estimate and parameter gradient at *θ*\* computed via reweighting using MBAR.

In Figure 4, we show this approach applied to retraining the ANI-2x model based on a training set of 212 tautomer pairs, where training is carried out until error began to increase on a validation set of 71 tautomer pairs. Figure 4**A** shows the learning curve, where the RMSE on the training set is reduced by over 4 kcal/mol. Surprisingly, there is almost no generalization gap (difference in between training and validation error), suggesting the information provided by the limited training set is sufficient to generalize to dissimilar tautomer pairs. Indeed, the test set error (evaluated on an independent set of 71 tautomer pairs) closely tracks the validation set error, and results in a model that has improved in RMSE from 7.2 [95% CI:6.1,8.5] kcal/mol to 3.2 [95% CI:2.7, 3.8] kcal/mol—an enormous increase in accuracy (Figure 4**B**).

**Figure 4.**
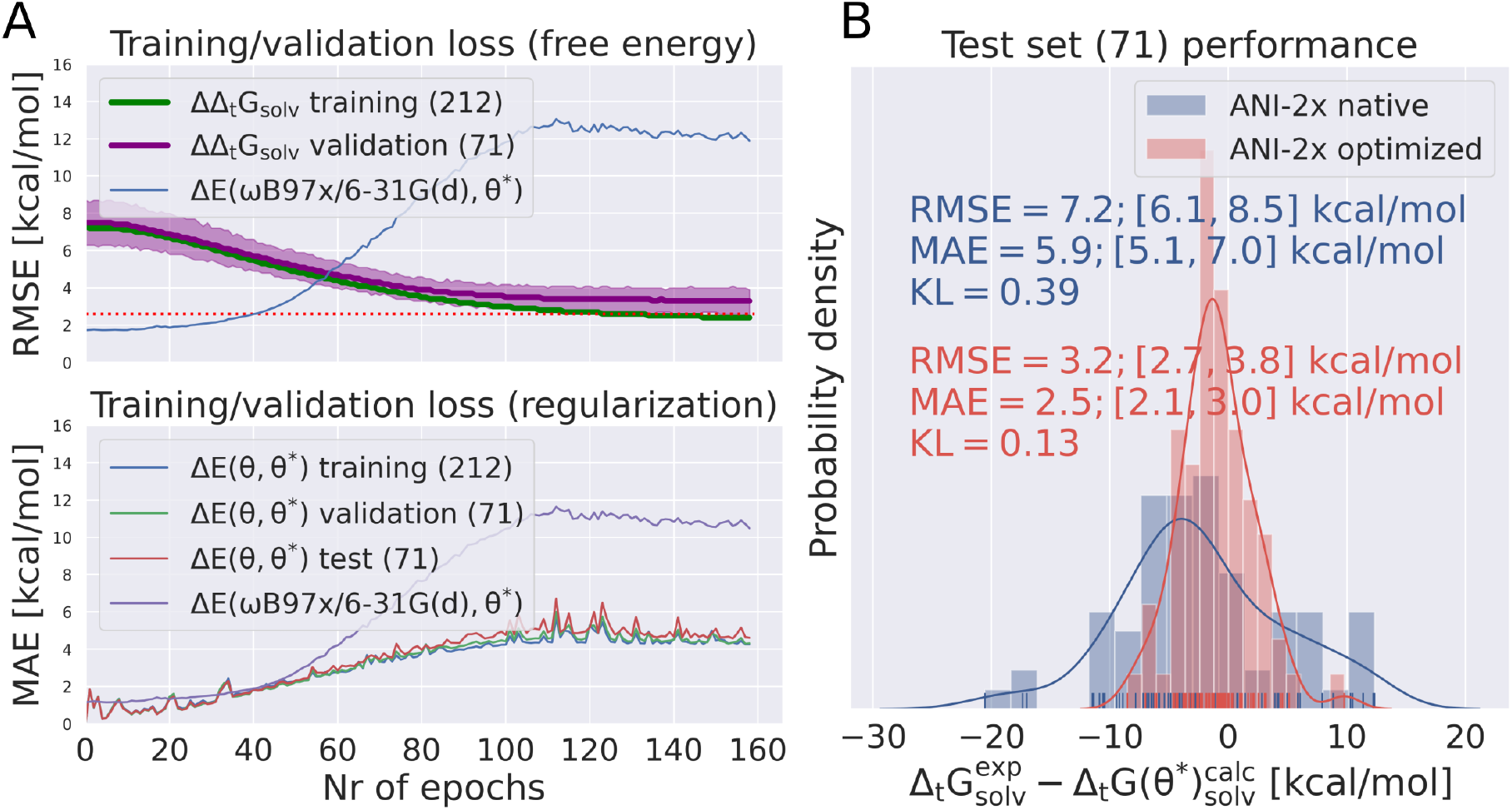
Quantum machine learning potentials can be optimized on a small set of experimental free energy training data to improve performance on an independent test set. The loss function for the retraining contained two terms: (1) the tautomeric free energy error (ΔΔ*_t_G_solv_*) and (2) the regularization error (Δ*E*(*θ, θ*\*)). In panel **A** the two terms of the loss function are plotted for the training/validation set as a function of the number of epochs. The single point energy error on the ANI-1x dataset (Δ*E*(*ω*B97x/6-31G(d), *θ*\*)) is shown in both plots, but was not part of the loss function and should only indicate the behavior of the new parameter set on the original ANI-1x training set. The top panel shows the RMSE of the training/validation free energy error and the bottom panel the regularization error. Panel **B** shows the distribution of the error of the estimated free energy with the original and the final ANI-2x parameters on a test set. The RMSE was reduced from initial 7.2 to 3.2 kcal/mol (the MAE was reduced from 5.9 to 2.5 kcal/mol).

Unsurprisingly, the improvement of the performance of the QML potential for the task of tautomeric free energy calculations leads to a decline in its accuracy on the ANI-1x QM dataset [50] (shown as ΔE(*ω*B97x/6-31G*(d), *θ*\*) in Figure 4 A, bottom panel). Ideally, this deviation would be included in the loss function and kept at a minimum. While we would recommend to include ΔE(*ω*B97x/6-31G*(d), *θ*\*) in the loss function for further, similar workflows we decided to omit it for technical reasons: evaluating the loss function was already very slow (for a ML application;very fast for a free energy calculation) and adding this particular penalty would have increased the time needed by roughly a factor of 5.

## Regularizing retraining by penalizing QML potential deviations does not degrade the ability to improve free energy predictions

To minimize the ANI-2x parameter set deviation from their original values a regularization term was added to the molecular loss function. This regularization term is *not* applied to deviation in parameter space, but is instead applied to deviation in the single point energy prediction these parameters imply on specific conformers. The regularization term is added to the molecular loss function and calculated as the mean absolute error (MAE) between energies computed using the the original parameter set *θ* and the perturbed parameter set *θ*\*, for all 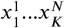 samples evaluated with the physical tautomeric endstate potentials *U*_1_ and *U_K_* (*K* represents the state and *N* the number of samples obtained from state *K*). This will be written as 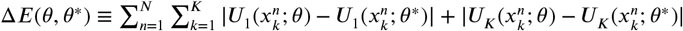. Evaluating the loss function for a single batch of the tautomer training set the regularization term calculated the energy deviation for 2 (number of physical endpoint potentials) × 11 (number of *λ* states) × 200 (snapshots per alchemical *λ* states) × 212 (training set tautomer pairs) resulting in a total of 932,800 data points. This regularization term does not hinder the ability to fit the free energy target if a reasonable weight is used for its contribution to the molecular loss function. It is difficult to compare the two terms in the loss function directly, since the free energy error (ΔΔ_*t*_*G_solv_*) is included as mean *squared* error and the single point energy (Δ*E*(*θ, θ*\*)) deviation as mean *absolute* error, but it seems that good results are achieved if the single point energy error scaling factor is below four times the scaling factor of the free energy scaling factor (the scaling factors used to generate Figure 4 are shown in Figure S.I.2). While retraining without the regularization term leads to equivalent improvements in tautomer free energy prediction accuracy, it produces huge deviations in the QML potential energies Δ*E*(*θ, θ*\*), as well as much larger deviations from the original ANI-1x dataset than retraining with the regularization term (Figure S.I.3).

There are two notable observations from the retraining without regularization: the large error on Δ*E*(*θ, θ*\*) does neither (1) impact its performance on the tautomer test set nor (2) translate to an error of similar magnitude on the ANI-1x dataset. We speculate that the reason for (1) is that the loss function optimizes relative free energy differences, allowing the absolute values to drift freely. This can be shown by plotting the magnitude of Δ*E*(*θ,θ*\*) with the physical endpoint potentials *U*_0_ and *U*_1_ on all samples 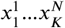 during the retraining. In Figure S.I.4 **A** we show this for a retraining with a loss function including a regularization term and in Figure S.I.4 **B** without a regularization term at four epochs (1,2,5,144/158). The Δ*E*(*θ, θ*\*) values support the hypothesis of shifted absolute values and are in agreement with the error of Δ*E*(*θ, θ*\*) for the full training/validation/test set in Figure S.I.3 **A**.

A possible explanation for (2) is the difference in the number of atoms of the systems that are compared. Δ*E*(*θ,θ*\*) indicates the difference in the absolute energy value of molecular systems with 220 to 250 atoms, multiplying small changes in the atom net for e.g. oxygen. This does not need to impact Δ*E*(*ω*B97x/6-31G(d),*θ*\*)) on the ANI-1x dataset similarly, even though both error terms seem to linearly increase through the optimization (but with different slope) as shown in Figure S.I.3.

The results shown in Figure 4 **B** highlight the incredible potential such a workflow has for optimizing QML parameters based on ensemble properties. The RMSE on the independent test set decreases from initial 7.2 kcal/mol to 3.2 kcal/mol.

## Retraining gradually increases the statistical error in reweighted free energies

As the QML model parameters *θ*\* deviate further from the initial model *θ*, the information about the equilibrium configuration probability density of *θ*\* provided by the equilibrium samples harvested at *θ* will diminish, and the statistical error will increase as the effective sample size decreases [51]. We examined how rapidly this occurs in Figure S.I.5, which depicts the statistical error in predicted free energy differences at epochs 40, 80, and 158 (the epoch as which training was stopped due to an increase in the validation set error). Surprisingly, the statistical error never exceeds 1.4 *k_B_T* for any system, despite the large number of training epochs carried out without any re-simulation to generate new equilibrium samples.

Interestingly, the rate at which different systems accumulate statistical error with parameter update epoch varies widely, with some systems barely increasing in statistical error. Using a cutoff of 1 *k_B_T* as acceptable for the perturbed free energy uncertainty, for example, Figure S.I.5 indicates that, for this system, only a few simulations would need to be re-run to generate updated equilibrium samples with the new parameter set *θ*\* after around 90 epochs to reduce their statistical error; a statistical error tolerance of 1.5 *k_B_T* would not require any re-simulation.

## Discussion

Quantum machine learning (QML) potentials have now come of age, providing a way to compute energies and gradients of molecular systems with orders of magnitude less computing effort than quantum chemical calculations. While current software and hardware implementations practically limit these simulations to small QML systems (or subsystems, as in [52]) at the moment, ongoing work in developing optimized hardware-accelerated software kernels to rapidly compute atomic feature vectors and continual improvements in hardware (such as GPUs) will rapidly increase the size and speed of these calculations, enabling them to be practical for treating entire solvated biomolecular systems within the next few years.

Given the rapid pace of software and hardware advances, alchemical free energy calculations with QML potentials offer multiple advantages over traditional MM potentials, as we have seen here: First, it is simple to implement alchemical free energy calculations, as the lack of singularities in QML potentials enables simple linear interpolations of physical endstate potentials to be used to compute statistically precise free energy differences. Second, QML potentials are suitable for treating phenomena that involve the breaking and making of bonds, such as free energies between tautomeric states, as we have shown here. Third, these potentials can be readily retrained based on a limited amount of experimental data in a manner that readily generalizes to improve predicted free energies on unrelated molecules. This retraining process can be rapidly carried out using reweighting techniques that avoid the need to frequently re-simulate. A regularization term included in the loss function can penalize deviations in the predicted quantum chemical energies that likely result from the accumulation of absolute energy offsets—to which free energy differences are insensitive—enabling the retraining process to be minimally perturbative to the original QML potential. Retraining only the final stages of the QML neural network, as we have done here, can be a practical choice that preserves many of the learned features in earlier stages of the network.

While the statistical error of estimated free energy differences does increase with each epoch of parameter optimization exclusively using reweighting, it appears to do so slowly, with none of the uncertainties exceeding 1.4 *k_B_T* (from an initial maximum of 0.7 *k_B_T*) during the 158 epochs of training required to produce the best model here. If great statistical precision is desired, re-simulating systems as this statistical error reaches a threshold could ensure all free energies remain within a desired level of precision throughout the retraining process. Because the *rate* at which the statistical error increased for each system varied greatly among tautomer pairs, it seems likely that only a small fraction of systems would require re-simulation at any one time, and this could be performed asynchronously to strike an optimal balance between controlling statistical error and reducing the wall clock time for re-optimizing the QML potential to experimental data.

Perhaps unsurprisingly, we found that QML potentials trained to higher levels of theory, such as the ANI-1ccx model, can produce better estimates of experimental tautomer free energy differences without retraining. Surprisingly, however, these highly accurate potentials can contain defects—such as the tendency for water molecules to become trapped in highly unusual geometries—that render them unsuitable for condensed-phase simulations. Care must be taken to either identify pathological configurations and abolish them through retraining in an iterative fashion (as appears to have been the case for ANI-2x) or physical asymptotic behavior must be built into the model to avoid this.

The ANI models provide an ensemble of neural networks whose deviation provides an estimate of the uncertainty of the QML potential in predicting the true target quantum chemical potential for the level of theory the model was parameterized for. This deviation has been used to target quantum chemical calculations to the most informative molecules using active learning approaches to minimize computational efforts in producing high-accuracy QML potentials [53]. Similar ideas could be used for designating experimental measurements that would be most useful in reducing the model uncertainty (as determined via reweighting to assess the variation in predicted free energies over individual neural network ensemble members) by targeting the collection of new experimental measurements using the deviation in predicted free energies over models as a target for uncertainty reduction.

The small generalization gap between test set performance and performance on both the validation and test sets is remarkable: often, machine learning approaches produce very large generalization gaps between training and validation sets. The small gap here suggests that the combination of retraining only the final layers of the QML potential neural network and the summed contribution of parameter gradients from many thousands of sampled configurations may serve to direct the model in highly generalizable directions. While we have not extensively assessed the data efficiency of this approach—how performance on held-out test data depends on the quantity of training data used—in detail, nor how designing test sets with chemical features similar or dissimilar to the test set impacts performance, it will be no doubt illuminating to study these properties in detail as the cost of these calculations continues to decrease due to advances in software and hardware.

## Code and data availability

- Python package used in this work (release v0.3): https://github.com/choderalab/neutromeratio
- Data to reproduce the plots/figures (release v1.0): https://zenodo.org/record/5245934

## Author Contributions

Conceptualization: JDC, JF, and MW; Methodology: JDC, JF, and MW; Software: JF, and MW; Investigation: JF, and MW; Writing-Original Draft: JDC, JF, and MW; Writing-Review&Editing: JDC, JF, and MW; Funding Acquisition: JDC and MW; Resources: JDC and MW; Supervision: JDC and MW.

## Acknowledgments

MW,JF and JDC are grateful for discussions within the Tautomer Consortium, specifically Paul Czodrowski, Brian Radak, Woody Sherman, David Mobley, Christopher Bayly, and Stefan Kast, as well as input from Adrian Roitberg and Olexandr Isayev. This work was only possible thanks to the time and effort Thomas Sander and Oya Wahl invested in curating the open tautomer database (Tautobase). MW is grateful for the support of Thierry Langer for use of computational resources and Oliver Wieder for helpful discussions.

## Funding

JF acknowledges support from NSF CHE-1738979 and the Sloan Kettering Institute. MW acknowledges support from a FWF Erwin Schrödinger Postdoctoral Fellowship J 4245-N28. JDC acknowledges support from NIH grant P30 CA008748, NIH grant R01 GM121505, NIH grant R01 GM132386, and the Sloan Kettering Institute.

## Disclosures

JDC is a current member of the Scientific Advisory Board of OpenEye Scientific Software, Redesign Science, and Interline Therapeutics, and has equity interests in Redesign Science and Interline Therapeutics. The Chodera laboratory receives or has received funding from multiple sources, including the National Institutes of Health, the National Science Foundation, the Parker Institute for Cancer Immunotherapy, Relay Therapeutics, Entasis Therapeutics, Silicon Therapeutics, EMD Serono (Merck KGaA), AstraZeneca, Vir Biotechnology, Bayer, XtalPi, Interline Therapeutics, the Molecular Sciences Software Institute, the Starr Cancer Consortium, the Open Force Field Consortium, Cycle for Survival, a Louis V. Gerstner Young Investigator Award, and the Sloan Kettering Institute. A complete funding history for the Chodera lab can be found at http://choderalab.org/funding.

## Detailed methods

### Experimental data

The Tautobase dataset [45] used in this study was obtained as a DataWarrior file deposited from https://github.com/WahlOya/Tautobase (commit 2b59c60 from of Jul 23, 2019). Tautobase is a machine-readable curated dataset of tautomer pair data, originally sourced from the Tautomer Codex authored by P.W. Kenny and P.J. Taylor [54]. From the dataset a subset of tautomer pairs were considered that

1. were measured/calculated/estimated in aqueous solution
2. had a numeric logK value between +/-10
3. had no charged species
4. did not contain iodine
5. only a single hydrogen and double bond change its position

476 of the 1680 deposited tautomer pairs had these properties. We added two tautomer pairs from the SAMPL2 challenge (Tautomer pair 2A_2B and 4A_4B) [55]. The logK value was converted to free energies with 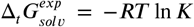. Further, tautomer pairs were removed that had incorrect molecular structures, stereobonds that changed its position or elements others than carbon, nitrogen, hydrogen and oxygen resulting in a final set of 354 tautomer pairs.

The subset of Tautobase used for the QML calculations can be found here as list of SMILES (354 tautomer pairs): https://github.com/choderalab/neutromeratio/blob/2abf29f03e5175a988503b5d6ceeee8ce5bfd4ad/data/ani_tautobase_subset.txt.

### Relative alchemical free energy calculations

Relative alchemical free energies were calculated using a single topology approach [5]. The hybrid topology (the superset of the two topologies) contained both the hydrogen being deleted and the hydrogen being added, differing by a single hydrogen from each of the physical tautomeric endstates.

By default, the coordinates of the hybrid topology were generated by using the coordinates of Tautomerl (as defined in the Tautobase database). If a cis/trans stereobond was created/removed in the tautomer reaction, the initial coordinates were taken from the topology with the stereobond present (therefore sometimes changing the direction of the tautomer reaction). The coordinates of the added, non-interacting (dummy) hydrogen were obtained by randomly sampling 100 positions on the surface of a sphere (with radius of 1.02 (Å), roughly corresponding to an X-H bond length) defined around its new bonded heavy atom, and subsequently selecting the position with the lowest ANI-2x energy. At the physical endstate, the dummy hydrogen interacted with the physical system only by a harmonic restraint keeping it in proximity to its bonded heavy atom (described in detail below). The molecule was solvated in a water droplet with a diameter of 16 Å around the center of mass. The alchemical free energy calculation used independent simulations at 11 equidistant states (two physical endstates with nine bridging alchemical intermediates).

Energies and forces were calculated using ANI-2x [28] as implemented in torchani https://github.com/aiqm/torchani). The alchemical system potential energy was defined by a linear combination of the initial (*U*_1_) and final (*U*_2_) tautomer potential energies as a function of the alchemical parameter *λin*[0,1]:

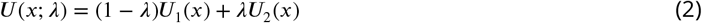

It is notable that, while simple linear combinations of physical endstates soft-core potentials and complex multistage protocols are generally required for alchemical free energy calculations with molecular mechanics potentials [56–58], the lack of singularities in quantum machine learning potentials like ANI-2x results in well-behaved, easily convergent calculations with simple linear interpolation to define alchemical potentials.

Before each simulation, initial coordinates were minimized using a BFGS optimizer as implemented in scipy [59]. Coordinates were sampled using Langevin dynamics at 300 K with a collision rate of 10 ps^-1^ and a 0.5 fs time step using the high-quality BAOAB integrator [60], which introduces little error in configuration space sampling [61]. Initial velocities were drawn from a Maxwell-Boltzmann distribution at the simulation temperature. 10,000 samples were obtained from 100 ps simulations at each alchemical state, saving coordinates each 10 fs.

Because nuclei can rearrange to form distinct chemical species in highly perturbed simulations, we applied a flat-bottom harmonic restraint to covalent bonds to ensure we sampled the desired chemical species during the free energy calculation. This restraint had the form:

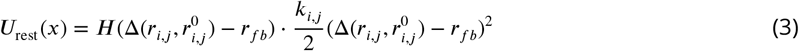

Here, *H* is the Heaviside step function, 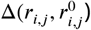 is the difference between the reference bond length 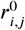 and the current bond length *r_i,j_* and *r_fb_* as half of the well radius. For all heavy atom pairs 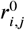 was set to 1.4 Å and *r_fb_* to 0.3 Å with *k_i,j_* set to *k_B_T*/0.1 Å. For C-H/O-H/N-H bond pairs 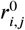 was set to 1.02 Å and *k_B_T*/0.2 Å and *r_fb_* to 0.4 Å. The restraint well was chosen so that the restraint does not interfere with normal bond stretching but will activate once a bond is broken.

Additional flat bottom restraints centered at the origin with a spring constant *k_B_T*/0.1 Å, with harmonic walls beyond the 8 Å droplet radius, were applied to all oxygen atoms of all water molecules. The center of mass of the molecule was restrained to the center of the droplet with a flat bottom restraint with a well radius of 0.1 Å. The same set of restraints was used for all lambda states of the alchemical sampling state for a tautomer pair but free energies were calculated without the restraints.

Relative alchemical free energy estimates 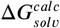 were calculated using the multistate Bennett acceptance ratio (MBAR) method as implemented in the pymbar package [51]. 300 uncorrelated snapshots were sampled from each alchemical state for use in MBAR analysis.

### Neural net parameter optimization based on experimental relative solvation free energies

#### Training/validation/testing split

The tautomer data set was randomly split (60:20:20) into training (212 tautomer pairs), validation (71), and test (71) sets.

#### Optimization and model selection details

Neural net parameters were optimized using a routine modified from the TorchANI tutorial [62]. To limit the capacity for overfitting, only the weights and biases of the *final* layer of each of the 8 pretrained ANI-2x models were optimized for each of the atom nets (one net per element), resulting in roughly 8×4×140 = 4480 tunable parameters. (ANI-2x is defined as an ensemble of 8 models, each containing one neural network per element with 4 linear layers and 3 RelU activation functions with a single bias parameter. The total number of parameters for the ensemble is 8×4700 (only counting the parameters for hydrogen, carbon, oxygen and nitrogen [28].) Another way to conceptualize this approach is to envision freezing the nonlinear “features” computed by the previous layers, and refitting the affine weights and bias used to combine these features into the final contribution to the overall potential energy in the final layer.

As in the TorchANI tutorial, the weight matrices were updated using the Adam optimizer with decoupled weight decay (AdamW), and the bias vectors were updated using Stochastic Gradient Descent (SGD). The learning rate for SGD was set to 10^-9^ and for AdamW to 10^-6^. The training data was randomly partitioned in each epoch in mini-batches of 7 tautomer pairs (this size was chosen because it was the maximum number of system were were able to process in parallel on a 128 GB RAM machine), and gradient updates were performed for each mini-batch. The best model was chosen based on the RMSE on the validation set, and model performance on the test set (not used in training or model selection) was reported.

#### Regularization of potential energy changes

The model was trained by minimizing the mean squared error (MSE) loss between calculated and experimental relative tautomeric free energies. The free energy error was calculated as

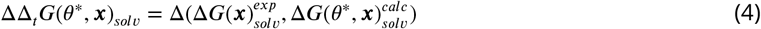

where *θ*\* denotes the perturbed parameter set and ***x*** denotes the conformations 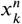, where 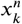 denotes conformation *n* samples from alchemical state *k*.

To avoid having the potential energies drift too far from true quantum chemical energies and minimize the opportunity to overfit the training set, we utilized a regularization term on computed tautomeric energy differences. While restraining to the original quantum chemical data used to fit the ANI-2x model [28] would have been optimal, this dataset has not been made public. Instead, we compute the regularization term as the mean absolute error (MAE) between the single point energy for the parameter set *θ*\* and the original parameter set *θ* on the set of conformations ***x***. The conformations ***x*** were evaluated using the potential energy functions at the physical end-states *U*_1_ and *U_k_*. In accordance with Figure 3, the regularization term for state 1 is

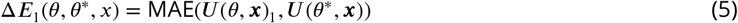

and the expression is analogous for U*_k_*. Both terms are then combined and a final term calculated with

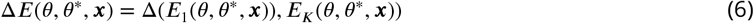

Δ*E*(*θ,θ*\*, ***x***) was normalized to better represent the contributions of tautomer systems with varying total numbers of atoms. The per-molecule pair (*m*) loss function *l* is defined as

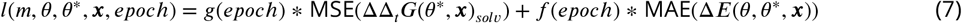

The two scaling factors *f*(epoch) and *g*(epoch) were used to control the contribution of the two terms as a function of training epoch. For the final results the values for *g*(labeled “scaling dG”) and *f*(labeled “scaling dE”) are shown in Figure S.I.2.

The overall loss is then

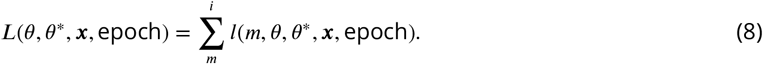

#### Free energy gradients by differentiable reweighting

The perturbed relative free energy difference 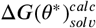 required for computing *l*(*m; θ*\*) is calculated by importance sampling, or “reweighting” the original MBAR estimate from the original parameters *θ* that were used during simulation, to the current parameter set *θ*\*. To compute this estimate efficiently at arbitrary *θ*\*, we first collect configuration samples ata reference value *θ* (corresponding to the original parameters of the pretrained ANI-2x model) for each intermediate value *λ*. For each configuration sample *x*, and each alchemical window *λ_k_*, we compute the reduced potential *u*(*x, λ; θ*), to form the *N*×*K* matrix of inputs to MBAR [40], where *N* is the total number of snapshots and *K* the number of alchemical states (*λ* windows).^1^ After the initial effort involved in simulating the samples *x^n^* and estimating the free energy differences among all alchemical states (*λ* windows), we can extrapolate free energy estimates (and their gradients with respect to parameters) to any nearby parameter set *θ*\* using the reduced potential *u*(*x, λ; θ*\*).

To do this, we note the log weight of sampled configuration *x^n^* in alchemical state *k* is given by log *q_k_*(*x^n^*) = *f_k_* – *u_k_*(*x^n^*). The MBAR estimator can be interpreted as constructing a mixture of all *k* states, with weights as a function of the number of samples *N_k_* available from state *k* and of the self-consistent free energy estimates *f_k_*. The total log weight of a sample *x_n_* within this mixture distribution is then given by

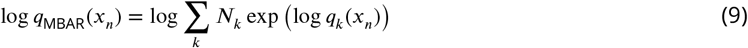

To estimate expectations in an arbitrary new state defined by unnormalized probability density *q*_new_(*x*) = *e*^-*u*_new_(*x_n_*)^, we compute importance weights *w*(*x_n_*).

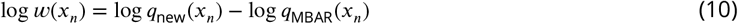

In particular, we can estimate the free energy in a new state *q*_new_ = *e*^-*u*_new_^ by

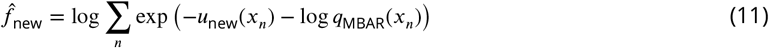

Thus, to compute relative free energies at a new value of the parameters *θ*\*, we need only compute *u*(*x, λ* = 1; *θ*\*) and *u*(*x, λ* = 0; *θ*\*) for all configurations *x*:

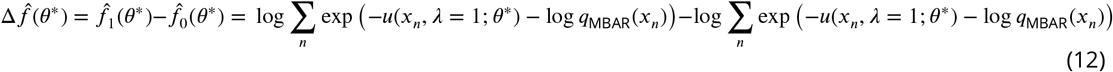

Assuming that *u*(*x, λ, θ*) is differentiable with respect to *θ*, the overall estimate is also differentiable with respect to *θ*.

Note that, if *θ*\* drifts sufficiently far enough from the original *θ* so that the statistical overlap between the new physical endstates and the sampled alchemical states is poor, we would need to initiate new simulations at the updated parameter set *θ*\*. To ensure this is not the case, the statistical error in our estimate (and the effective sample size) were closely monitored during the optimization process. Reported estimates of tautomer free energies include estimates of the statistical error in the MBAR-extrapolated free energies.

## Supplementary Information

### Analysis of ANI-1ccx droplet simulation

We decided to focus on ANI-2x instead of ANI-1ccx because of preliminary results obtained by simulating a small molecule in a 16Å water droplets with ANI-1ccx. All observed ANI-1ccx droplet simulations revealed nonphysical H-O-H minimum angle geometries that we were not able to avoid or control (H-O-H angles during the course of a 200 ps simulation with ANI-1ccx are shown in Figure S.I.1 **A**). These nonphysical water geometries had two interesting effects: (1) water molecules adopting these geometries contributed highly favorable to the total energy of the system (shown for the water oxygen in Figure S.I.1 **B**) and (2) the ensemble standard deviation (the standard deviation calculated for the per atom energy prediction of the 8 neural networks) increased significantly for water molecules adopting these geometries (shown for the water oxygen in Figure S.I.1 **C** and **D**). While we were able to perform relative alchemical free energy calculations with molecular systems containing these nonphysical water geometries (their geometry were not a result of the alchemical part of the potential and did not dependent on *λ*) its impact on the results was not quantifiable. We show the results for the alchemical free energy calculations with ANI-1ccx in Figure 2 as well as detailed plots for the nonphysical water angles in Figure S.I.1.

**Appendix 0 Figure S.I.1.**
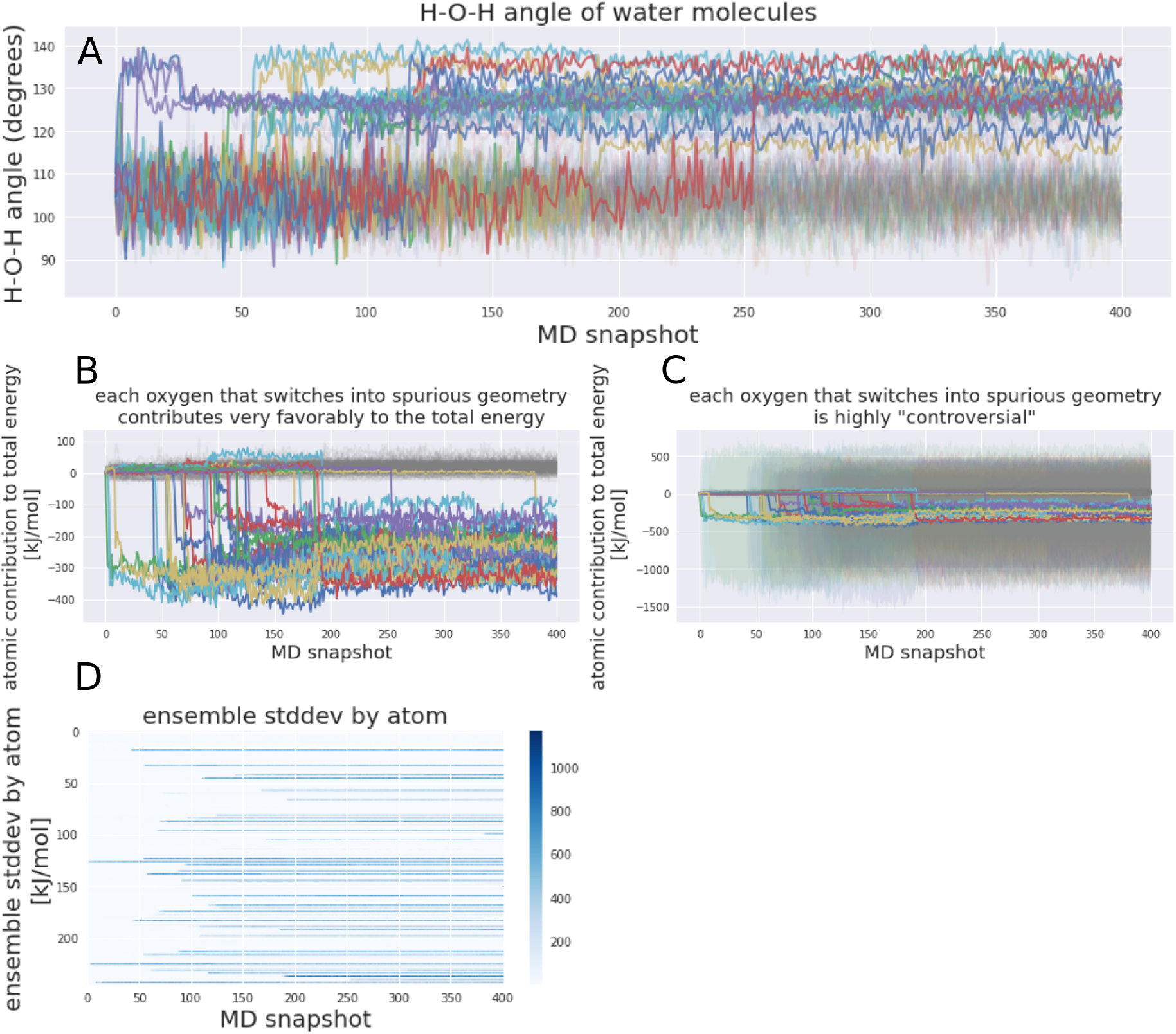
ANI-1ccx water droplet simulations show nonphysical geometry in the H-O-H angle of water molecules. Results are plotted for a 200 ps water droplet simulation. In panel **A** the water H-O-H angles are plotted for each water molecule during the simulation. We observe nonphysical H-O-H angle (other than 104 degrees) nearly immediately after the simulation starts for an increasing number of water molecules. These water geometries also seem to be highly stable and no transition from the nonphysical geometry to the physical H-O-H angle was observed. Panel **B** plots the atomic contribution of oxygen atoms for each water molecule and shows that oxygen atoms of nonphysical water contribute significantly to the total energy of the system. Panel **C** plots again the atomic contribution of oxygen atoms to the total energy (as Panel **B**) but adds the ensemble standard deviation for each water molecule adopting non-physical geometries. The magnitude of the standard deviation is in the order of 1,000 kJ/mol. Panel **D** plots again the ensemble standard deviation for each oxygen atom.

**Appendix 0 Figure S.I.2.**
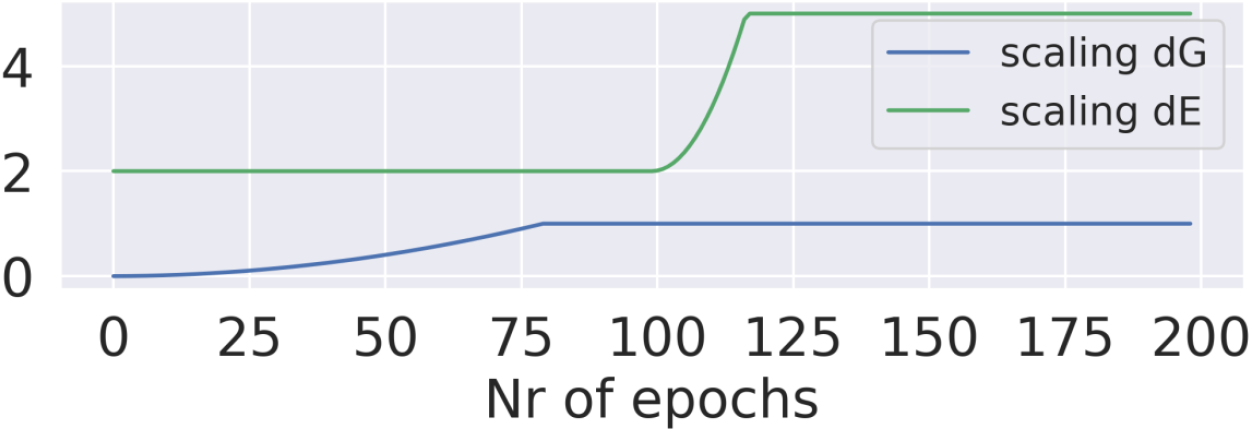
The scaling factors for f(epoch) and g(epoch) used in the molecular loss function in the reported results in 4.

**Appendix 0 Figure S.I.3.**
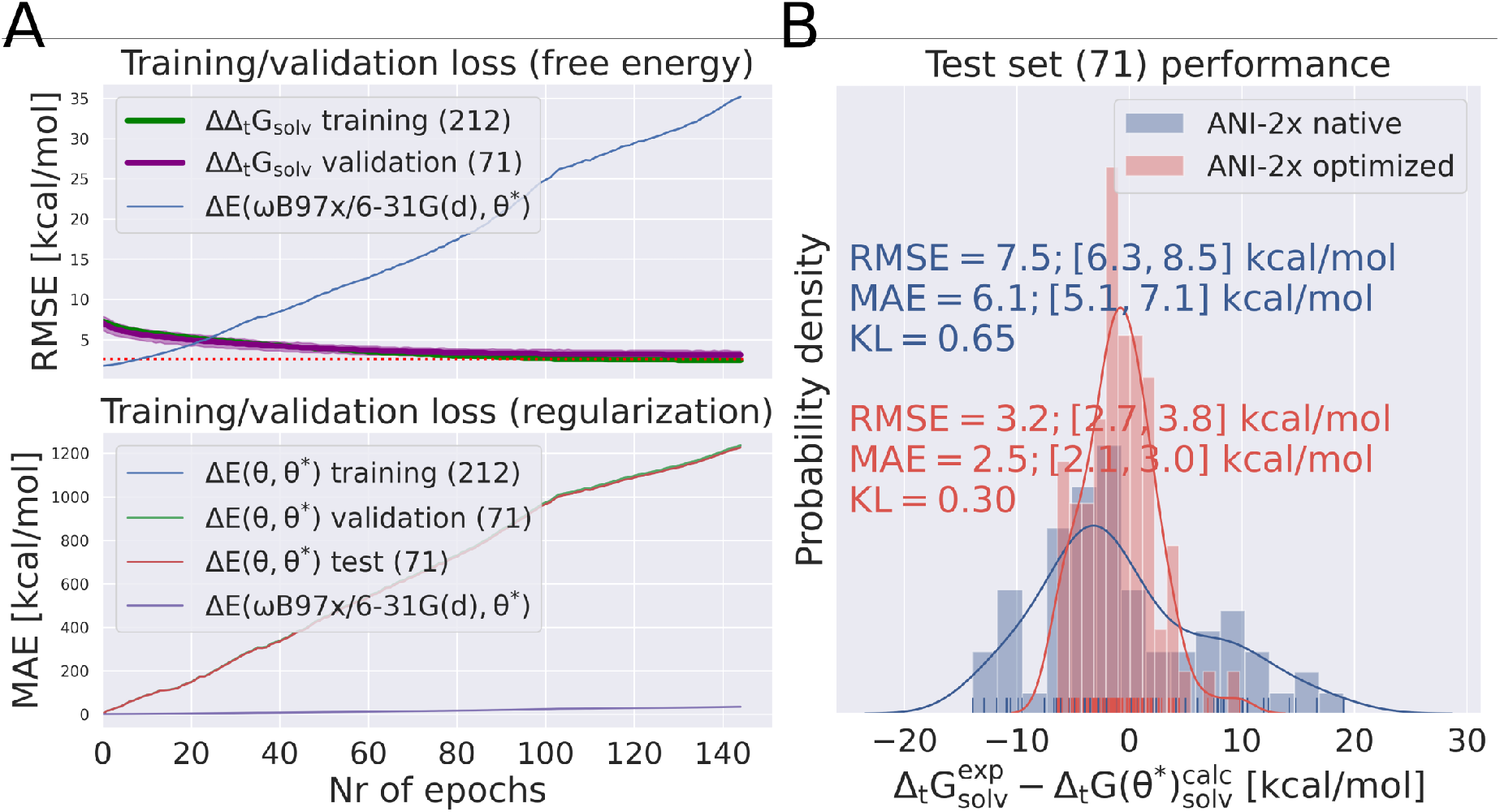
The training/validation loss and test set performance is shown for a retraining run without applying the regularization term in the loss function. The top panel shows the RMSE of the training/validation free energy error (Δ*_t_G_solv_*) for the training/validation set and the bottom panel the MAE of regularization error (Δ*E*(*θ, θ*\*)) (which is *not* included in the loss function). The single point energy error on the ANI-1x dataset (Δ*E*(*ω*B97x/6-31G(d), *θ*\*)) is shown in both plots, but was not part of the loss function and should only indicate the behavior of the new parameter set on the original ANI-1x training set. Panel **B** shows the distribution of the error of the estimated free energy with the original and the final ANI-2x parameters on a test set. The RMSE was reduced from initial 7.5 to 3.4 kcal/mol (the MAE was reduced from 6.1 to 2.6 kcal/mol)

**Appendix 0 Figure S.I.4.**
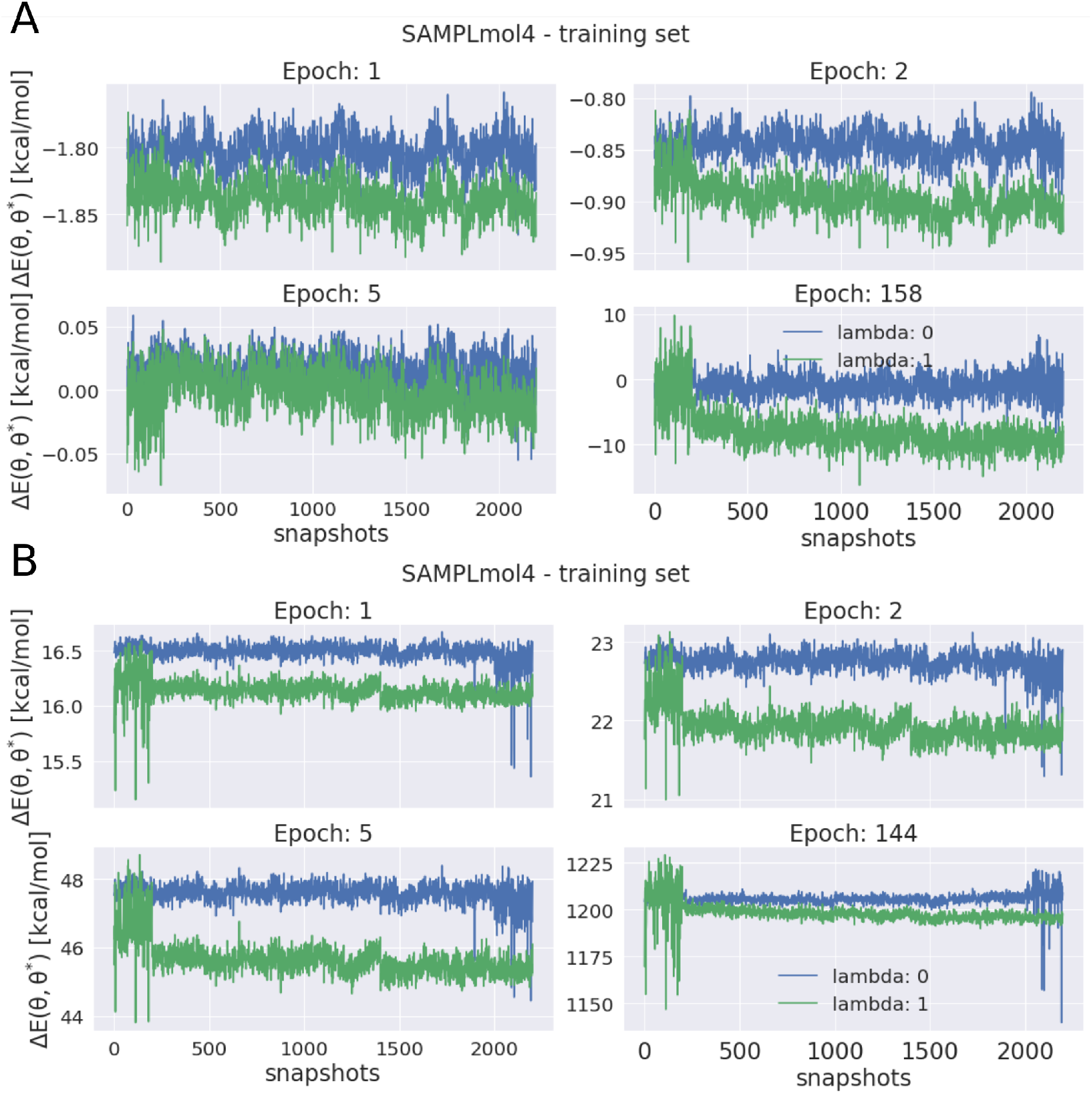
Regularization restricts the shifting of energy offsets during retraining. Δ*E*(*θ,θ*\*) is shown for the endpoint potentials on samples 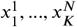 at different epochs for a single tautomer pair from the training set. **A** shows plots for a retraining run with regularization, **B** without regularization. The magnitude of Δ*E*(*θ,θ*\*) for the 4 epochs shown in **A** is relatively similar compared to the 4 plots in **B**, in which Δ*E*(*θ, θ*\*) steadily increases.

**Appendix 0 Figure S.I.5.**
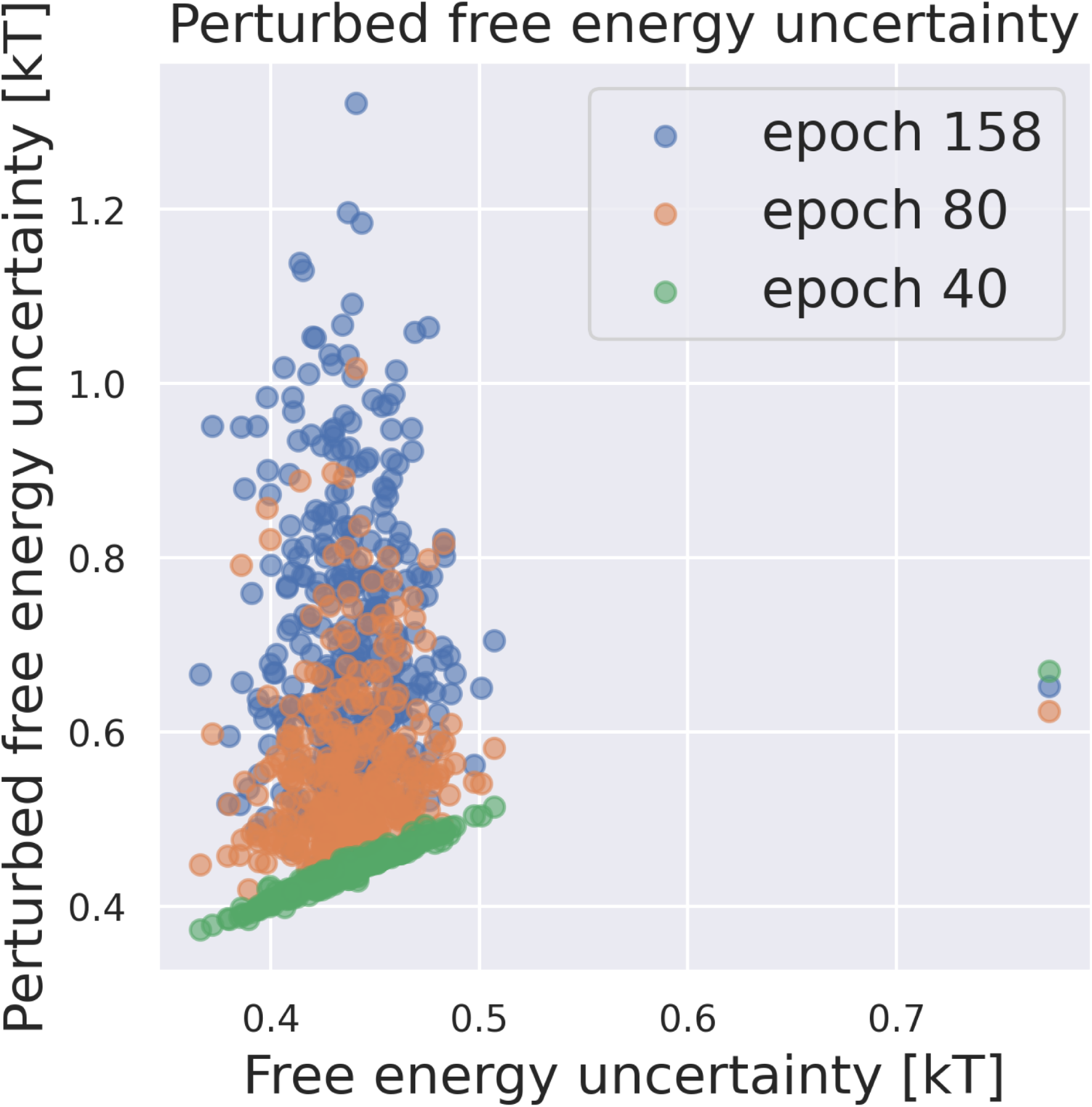
Perturbed free energy uncertainty increases with the number of training epochs. The free energy estimate uncertainty was calculated using the MBAR implementation in pyMBAR. The free energy uncertainty before retraining is shown on the x-axis.

## Footnotes

1 Here, 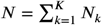, with *N_k_* = 200 uncorrelated snapshots per *λ_k_* state used for the MBAR estimate.

## Notes

### Competing Interest Statement

The authors have declared no competing interest.

### Summary of Updates

Adding paragraph to introduce the importance sampling approach in the introduction and highlight how samples from a distribution can be "reweighted" to obtain a different target distribution.

